# Limits of optimal decoding under synaptic coarse-tuning

**DOI:** 10.64898/2026.02.10.705038

**Authors:** Ori Hendler, Ronen Segev, Maoz Shamir

## Abstract

Sensory information propagates through successive processing stages in the brain, where synaptic weight patterns between stations determine how downstream neurons decode information from upstream populations. Although optimized synaptic connectivity can enhance information transmission, it requires precise weight tuning. Recent evidence depicting substantial synaptic volatility raises two fundamental questions: How does coarse-tuning of synaptic connectivity affect information transmission? What strategies could the nervous system employ to maintain reliable communication despite synaptic fluctuations? We addressed these questions by analyzing the signal-to-noise ratio (*SNR*) for binary stimulus discrimination under two decoding schemes: a naïve population average and an optimized linear decoder. For the naïve decoder, we found that *SNR* remains largely insensitive to synaptic imprecision, since performance is already limited by correlated noise in neuronal responses. For the optimal decoder, we identified three distinct regimes, that is, weak, moderate and strong coarse-tuning. Under weak coarse-tuning, *SNR*^2^ scales linearly with population size *N*. Under moderate coarse-tuning, scaling becomes sublinear. Strikingly, under strong coarse-tuning, the regime most consistent with observed neuronal heterogeneity, *SNR* saturates and can not be improved by recruiting larger populations. This limitation persists even when incorporating feedforward or recurrent network architectures. These findings suggest that in the biologically relevant regime of strong coarse-tuning, naïve and optimal decoders can achieve qualitatively similar performance. The analysis shows that effective readout under synaptic volatility is constrained to an invariant low-dimensional manifold aligned with the naïve decoder, potentially pointing to a fundamental principle for robust neural computation in the face of ongoing synaptic remodeling.

## I. INTRODUCTION

One of the main enigmas in neuroscience is how sensory information is transmitted and decoded as it propagates downstream [1]. The profile of synaptic weights largely determines the fidelity of information transfer along the processing pathway [2–4]. Accordingly, the synaptic-weight profile can be regarded as a decoder or a readout algorithm [5, 6]. One such possible decoder is a naïve readout that assigns uniform weights to all inputs. While easy to implement, it generally results in suboptimal performance [7–11]. By contrast, an optimal decoder can leverage detailed knowledge of the population response statistics to achieve superior performance by fine-tuning its synaptic weights to the detailed statistics of the network activity [12–15]. The caveat is that synaptic weights undergo substantial remodeling.

A growing body of empirical evidence has demonstrated that synapses are highly volatile [16, 17]. In-vivo studies show that dendritic spines, which reflect synaptic strength, exhibit substantial size fluctuations over time scales ranging from a few hours to days [18–21]. In-vitro studies of cultured cortical neurons show that synaptic sizes fluctuate dramatically at a time scale of hours [22–25], with considerable fluctuations on a similar scale emerging even in the absence of neuronal activity [26–Theoretical studies typically partition synaptic weights into structured and unstructured components [29– The structured component captures the tuned connectivity, whereas the unstructured component represents random variability, which we refer to as the coarse-tuning of the weights, which may or may not fluctuate over time [18, 34–40].

Considerable theoretical attention has been devoted to querying how the brain can maintain functionality in the face of considerable coarse-tuning of synaptic weights. To date, however, the computational implications of the coarse-tuning of synaptic weights on the fidelity of information transfer in the brain remain largely unknown.

Here, we investigated the effect of the coarse-tuning of synaptic weights on readout accuracy in the framework of a modeling study. We begin by defining a statistical model of neuronal responses to the stimulus. We then define two decoders: a naï ve decoder and an optimal one. We then apply coarse-tuning to the synaptic weights of both decoders, and investigate its effect on the decoding accuracy. Next, we consider two extensions of our readout model: a multilayer Feedforward Neural Network architecture and a Recurrent Neural Network. We show that neither can mitigate the drastic detrimental effect of coarse-tuning. The discussion centers on the implications of these findings for the theory of population coding in the brain.

## II. STATISTICAL MODEL OF THE NEURAL RESPONSE

We model a population of *N* neurons coding for a binary stimuli *α* ∈ {*t, d*} (target or distractor). We assume that the neural responses **r**, conditioned on stimulus *α*, is distributed according to multivariate Gaussian statistics with mean

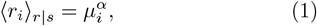

and covariance matrix

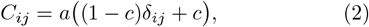

where ⟨·⟩ denotes averaging over neural responses for a given stimulus. The parameter *a* is the single neuron variance and *c* is the pairwise correlation coefficient. A Gaussian approximation offers a tractable and empirically-based description of neural responses [41]. The population is heterogeneous, where each neuron has distinct mean response parameter 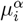 randomly drawn when the network is created and fixed thereafter. Trial-to-trial fluctuations thus arise from the noise structure whereas variability in 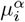 reflects frozen heterogeneity across neurons. Different network realizations yield different parameter sets ***µ***^*α*^, with first-order statistics

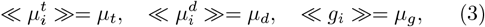

where double brackets, ≪ · ≫, denote ensemble average over realizations. The second-order statistics are

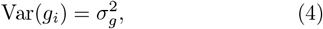

where we assume that 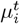 and 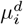 are uncorrelated and have equal variance 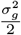.

We further assume that the neuronal responses exhibit stimulus selectivity, where on average, responses to the target exceed responses to the distractor. We quantify response selectivity as

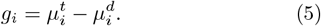

## III. THE READOUT MODEL

To examine the accuracy at which information about a stimulus can propagate downstream, we study readout performance in the framework of a two-interval-two-alternative forced-choice task. In this paradigm, one interval presents the target stimulus and the other the distractor, with their order randomized across trials. Without loss of generality, we assume that the target is presented in the first interval for all analytical calculations. The linear readout discriminates between the two alternatives according to the sign of the field:

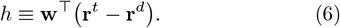

Thus, the linear readout is fully defined by its weight vector **w**. The binary decision follows from the sign of this field:

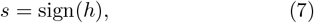

where *s* = +1 indicates “target first” (correct under our assumption) and *s* = −1 indicates “target second” (incorrect). In our Gaussian response model, the field *h* varies from trial to trial with mean ⟨*h*⟩ = **w**^⊤^ (***µ***^*t*^ − ***µ***^*d*^) and variance ⟨ (*δh*)^2^⟩ = 2 **w**^⊤^**Cw**, contingent on the specific realization of **w**. For a given realization, the probability of error is therefore

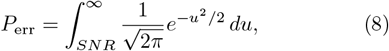

where the signal-to-noise ratio (*SNR*) squared is given by

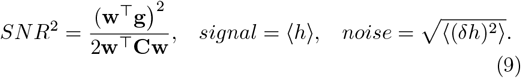

The probability of error is fully determined by the *SNR*. For large values of the *SNR*, a standard asymptotic expansion yields

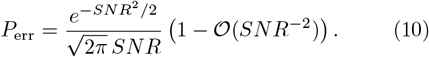

where the error probability decays exponentially with *SNR*^2^ up to algebraic corrections. Accordingly, we study how different choices of the weight vector **w** impact the *SNR* and hence task performance, see Fig.1a.

**FIG. 1:**
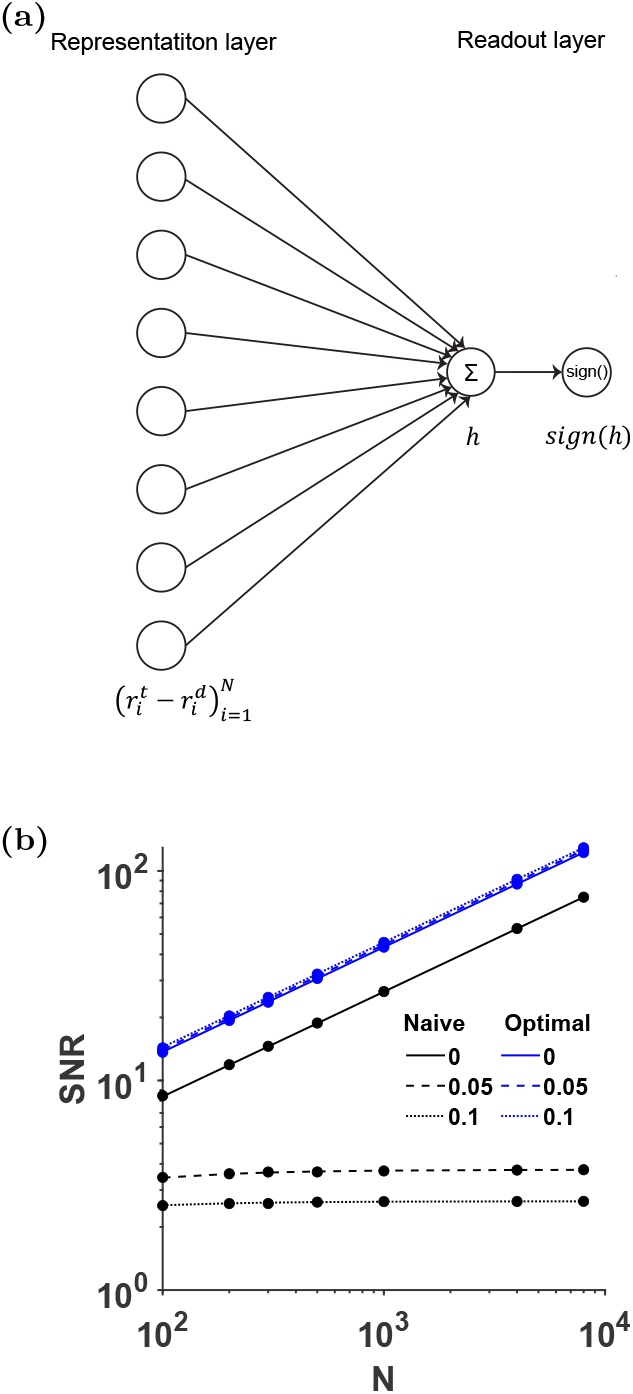
Readout model of the naïve and optimal decoders. **(a)** Schematic illustration of the model architecture. **(b)** The signal-to-noise ratio (*SNR*) is plotted as a function of population size *N* for the optimal (blue) and naïve (black) decoders. Different values of the pairwise correlation coefficient *c* = 0, 0.05, 0.1 are depicted by solid, dashed, and dotted lines, respectively.

## IV. READOUT PERFORMANCE

### A. Naïve Decoder Performance

We analyze the *SNR* as a function of different choices of readout weights, starting with the Naïve decoder. Since the response to the target is larger on average than the response to the distractor, the naïve approach consists of assigning equal weights to all neurons:

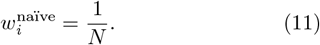

The *noise*^2^ (see Eq.(9)) of the naïve decoder is 𝒪(*N*^0^) and does not depend on the specific realization of quenched disorder, see Appendix A. The *signal*^2^ of the naïve decoder is a random variable that depends on the specific realization of the system; i.e., the choice of 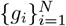. The *signal*^2^ has a mean of 𝒪(*N*^0^) and variance 𝒪(*N* ^−1^) across different realizations. Thus, for large populations (*N*→ ∞), the *signal*^2^ of a typical realization is equal to its quenched mean, up to fluctuations that vanish in the limit of large *N*. This property of a random variable whose fluctuations vanish relative to its mean is referred to as self-averaging. In the limit of large populations, a self-averaging variable can be replaced by its quenched mean. The quenched mean of the squared signal-to-noise ratio for the naïve decoder is (see Appendix A):

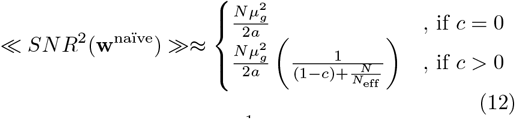

where we define 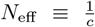. When response fluctuations of different neurons are uncorrelated, *c* = 0, the *SNR*^2^ scales linearly with the population size, *N*, ≪ *SNR*^2^(**w**^naïve^) ≫ ∝ *NI*_0_, where 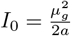 is the (quenched mean of the) *SNR*^2^ of the single neuron, Fig.1b (solid black line). For *c >* 0, the *SNR*^2^ scales linearly with the population size, ≪ *SNR*^2^(**w**^naïve^) ≫ ∝ *NI*_0_, for small populations, *N* ≪ *N*_*eff*_. In the limit of large populations, *N* ≫ *N*_*eff*_, the *SNR*^2^ of the naïve readout saturates to ≪ *SNR*^2^(**w**^naïve^) ≫ ∝ *N*_*eff*_ *I*_0_, Fig.1b. Note that due to self-averaging, the error bars denoting the standard deviation of the *SNR*^2^ across different realizations of the quenched disorder are barely visible.

### B. Optimal Decoder Performance

The optimal weights that maximize the probability of correct discrimination are given by:

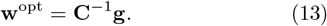

In our model; i.e., Eq. 2, this yields:

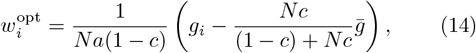

where 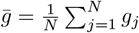 is the population-average response selectivity. In the limit of a large population (*N* → ∞) the weights are given by

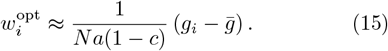

Hence, the optimal decoder uses the heterogeneity to extract information from a subspace that is orthogonal to the space with large noise correlations. We use this approximation, Eq. 15, for the analysis below. The *SNR*^2^ of the optimal decoder (Eq. (14)) is given by (see Appendix B):

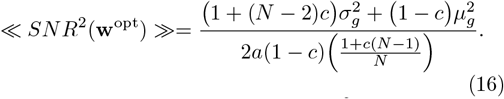

In a homogeneous population; i.e., 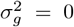, the optimal readout is the naïve readout and its *SNR* is limited by the noise correlations. In a heterogeneous popula-tion, 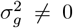, the *SNR* of the optimal readout is superior to that of the naïve one. In particular its *SNR*^2^ scales linearly with the population size, the leading term: 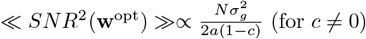, see Fig.1b (blue lines). However, these results rely on the ability of the system to finely tune the readout weights to the specific realization of the response heterogeneity; e.g., Eq. 14.

## V. COARSE-TUNED READOUT MODEL

Up to now, the quenched variability was only applied to the neuronal responses (i.e., 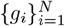, but not to the synaptic weights. To model synaptic volatility we introduce additive quenched fluctuations to the weights,

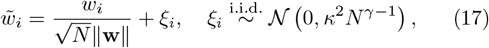

where the fluctuations *ξ*_*i*_ are independent and identically distributed Gaussian variables, with zero mean and variance 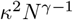. The parameter *κ* governs the magnitude of the fluctuation; i.e., the coarse-tuning magnitude. The parameter *γ* determines the scaling of the fluctuation variance with the population size, *N* ; i.e., the coarse-tuning scaling.

Thus, the weight 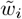 is expressed as the sum of a structured component of order 𝒪(*N* ^−1^) and an unstructured component that scales like 𝒪(*N*^*γ*^). This decomposition into structured and unstructured components is a classic feature of theoretical models, such as the Hopfield model [34], the Sherrington–Kirkpatrick (SK) model [42], and spin-glass models of neural networks [37, 43], as well as more recent works [29, 30, 44, 45]. In these models, the structured component of the weight vector **w** typically scales as 𝒪(*N* ^−1^), whereas the unstructured component scales as 𝒪(*N* ^−^ 2), which in our notation corresponds to *γ* = 0.

## VI. IMPACT OF COARSE-TUNING ON READOUT PERFORMANCE

The analysis of the effects of coarse-tuning on the *SNR* of the naïve and optimal readouts (below) identifies three regimes: weak (*γ* = −1), moderate (−1 < *γ* < 0) and strong (*γ* = 0) coarse-tuning. For weak coarse-tuning (*γ* = −1), the *signal* is self-averaging for both readouts,with mean 𝒪(*N*^0^) and variance of order 𝒪(*N* ^−1^), see Appendix C, D. Fig.2a presents the quenched distribution of the *signal* for weak coarse-tuning across the different realizations of response selectivity, 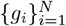, and weights, 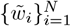, where each row corresponds to a different population size, *N*. As shown in the figure, the variability of the *signal* vanishes with increasing *N*. For weak coarse-tuning, the *noise*^2^ of the naïve decoder is self-averaging, Fig.2b (left column). However, the *noise*^2^ of the optimal decoder is not self-averaging, Fig.2b (right column), with mean 𝒪(*N* ^−1^) and variance 𝒪(*N* ^−2^). Nevertheless, for both decoders, the *SNR* is well-approximated by 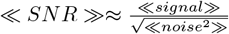, Fig.2c.

**FIG. 2:**
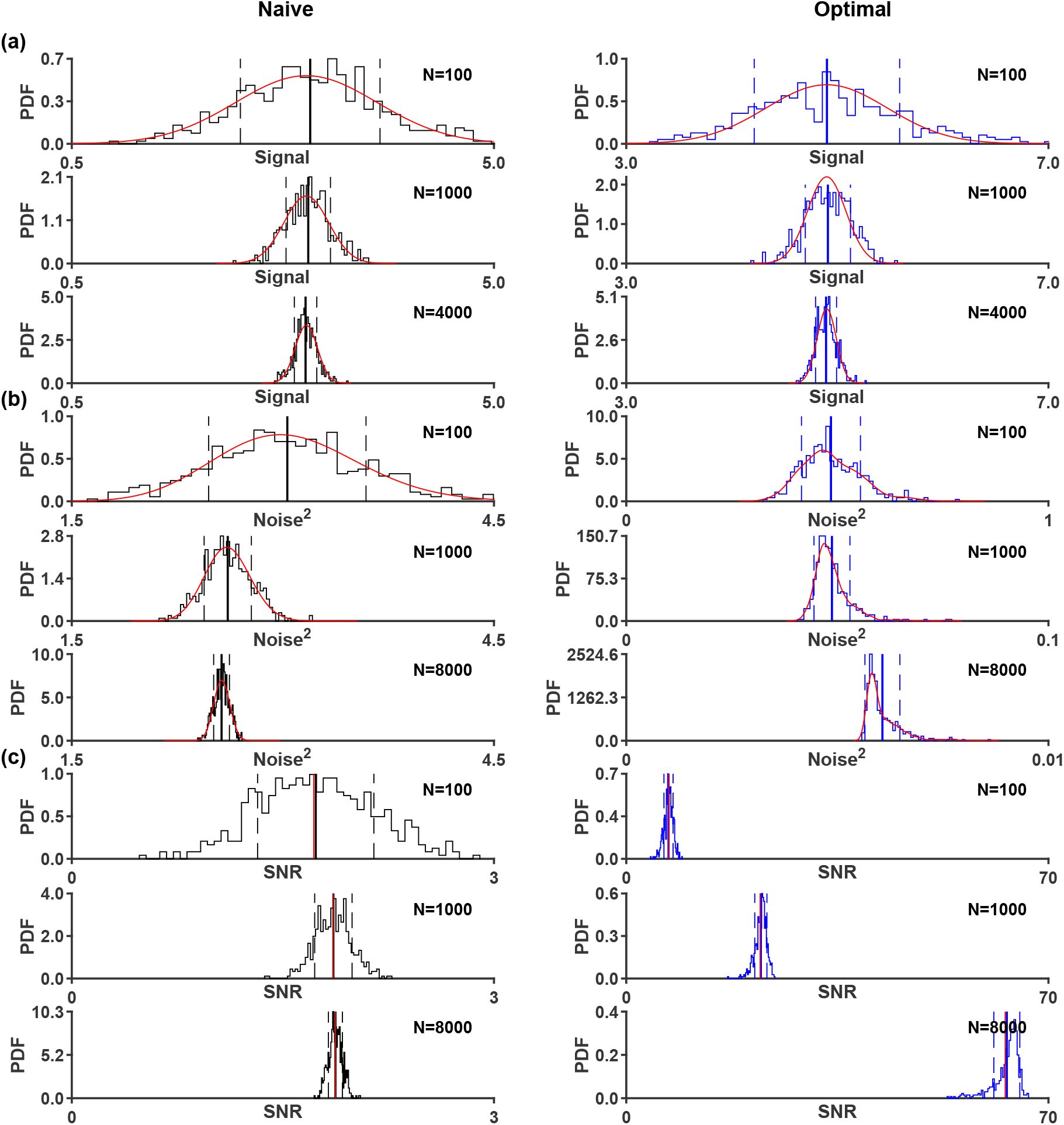
Histograms of signal, noise^2^, and SNR. Distributions of the naïve (black) and optimal (blue) decoders across population sizes *N* ∈ {100, 1000, 4000} under coarse-tuning with *κ* = 1 and *γ* = −1. **(a)** *signal* distribution. **(b)** *noise*^2^ distribution. **(c)** *SNR* distribution. Solid vertical lines indicate the mean; dashed lines indicate the ±1 standard deviation. Red curves show analytical predictions derived in the main text.

Similar behavior is found for moderate coarse-tuning. Specifically, the *signal*^2^ of both readouts is self-averaging. The *noise*^2^ is self-averaging for the naïve decoder (Appendix C), but not for the optimal decoder (Appendix D). In the strong coarse-tuning regime (*γ* = 0), neither the *signal* nor the *noise* are self-averaging. Nevertheless, ≪ *SNR* ≫ is well-approximated by the ratio of the averaged *signal* to the square root of the averaged *noise*^2^; i.e., 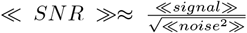. The quenched mean ≪ *SNR* ≫ provides a fair approximation of the typical *SNR*. We therefore use the *signal* and *noise*^2^ to quantify the *SNR* below.

We find that for large *N* the quenched mean of the *SNR* of the naïve and of the optimal readouts is well-approximated by (see Appendix C, D):

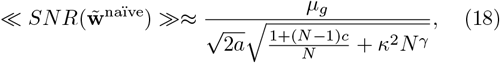

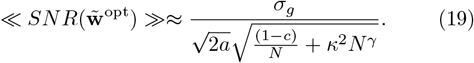

As shown in Eqs. (18)-(19) the nominator is independent of the population size and the scaling of the denominator with *N* depends on *γ*.

In the weak coarse-tuning regime, *γ* = −1, the *SNR*^2^ of the optimal decoder scales linearly with *N*. The slope of the linear scaling is determined by the magnitude of the quenched fluctuations, *κ* (see Fig.3a). For moderate coarse-tuning, −1 < *γ* < 0, the *SNR*^2^ grows with the population size, albeit in a sublinear manner, scaling as *N* ^−*γ*^, Fig.3b. For strong coarse-tuning, *γ* = 0, the *SNR* saturates to an asymptotic value,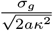, in the limit of large *N*, Fig.3b and Fig.3e.

**FIG. 3:**
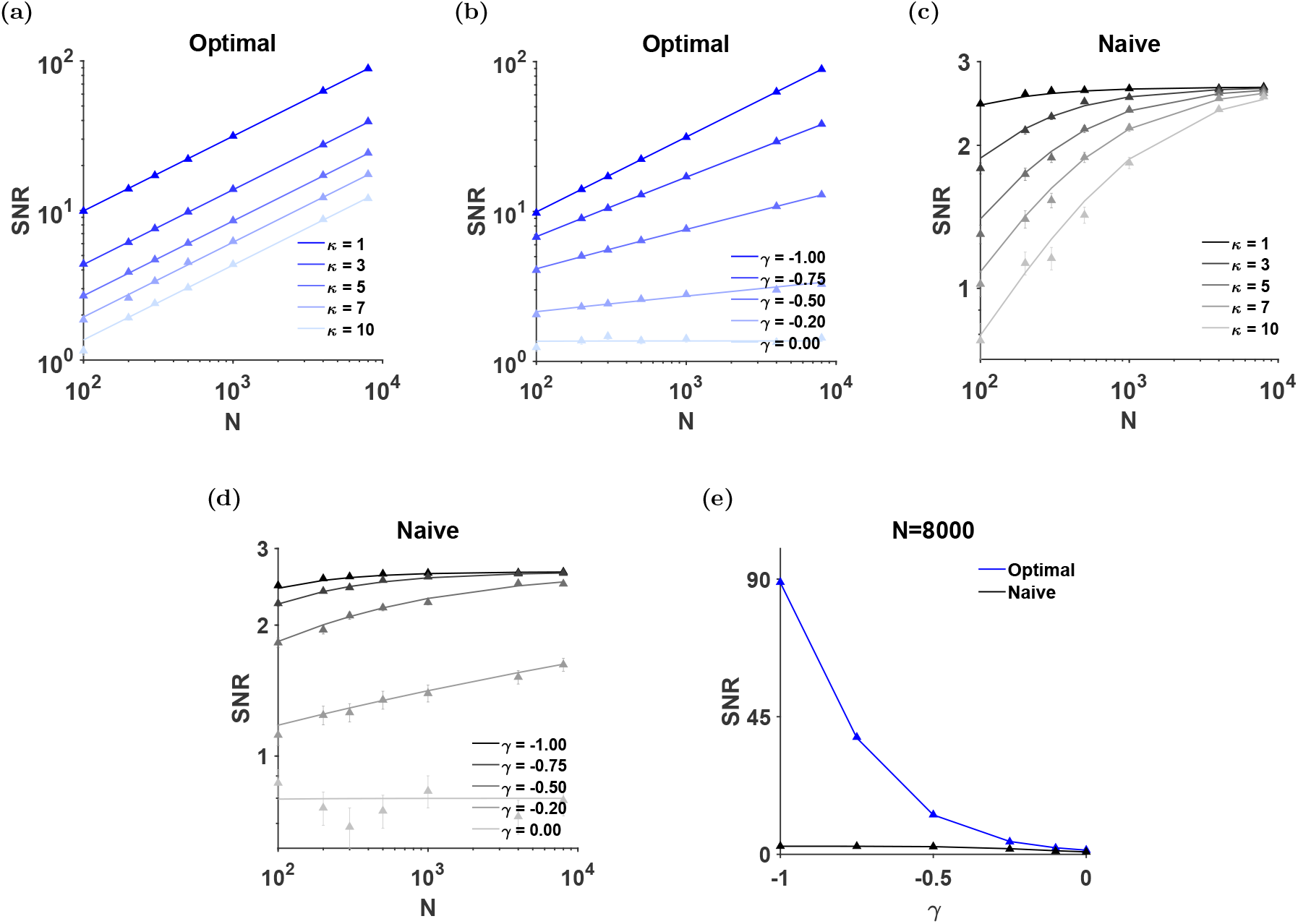
*SNR* scaling with population size under coarse-tuning. **(a)** *SNR* versus population size *N* for the optimal decoder, with varying magnitudes of coarse-tuning, *κ* ∈{1, 3, 5, 7, 10} at fixed *γ* =−1. **(b)** *SNR* versus *N* for the optimal decoder, with varying scalings of coarse-tuning, *γ* ∈{−1, −0.75, −0.5, −0.2, 0} at fixed *κ* = 1. **(c**,**d)** Same as (a,b), but for the naïve decoder. **(e)** *SNR* versus *γ* at fixed population size *N* = 8000 and *κ* = 1. In all panels, the color gradient from dark to light indicates increased coarse-tuning.

The *SNR* of the naïve decoder saturates in all coarsetuning regimes. For weak and moderate coarse-tuning,−1 ≤ *γ* < 0, the asymptotic value of the naïve *SNR* is determined by the noise correlations, 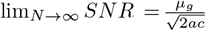, and is independent of the quenched disorder, Fig.3c and Fig.3d. In the case of strong coarse-tuning, *γ* = 0, the *SNR* of the naïve readout saturates to 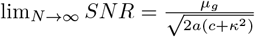, Fig.3e.

In summary, in weak and moderate levels of coarsetuning, −1 ≤ *γ* < 0, the optimal readout is superior to that of the naïve readout, and its *SNR* increases with the population size. In the strong coarse-tuning regime, *γ* = 0, the *SNR* of both readouts saturate to a finite limit. Moreover, in this case, the performance of the naïve readout can be superior to that of the optimal readout, depending on the choice of parameters. This raises the question of whether the performance of the optimal readout in the case of strong coarse-tuning can be improved by considering a more elaborate readout structure incorporating more layers and recurrent connectivity.

## VII. READOUT LAYER AND FURTHER PROCESSING

### A. Feedforward Neural Network

To incorporate an additional processing layer, we treat the binary decision variable *s* (Eq. 7) as a single neuron in the processing layer of *N* neurons, 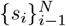 with *s*_*i*_ = sign(*h*_*i*_) (see Fig.4a), where *h*_*i*_ is the local field of neuron *i* and is given by

**FIG. 4:**
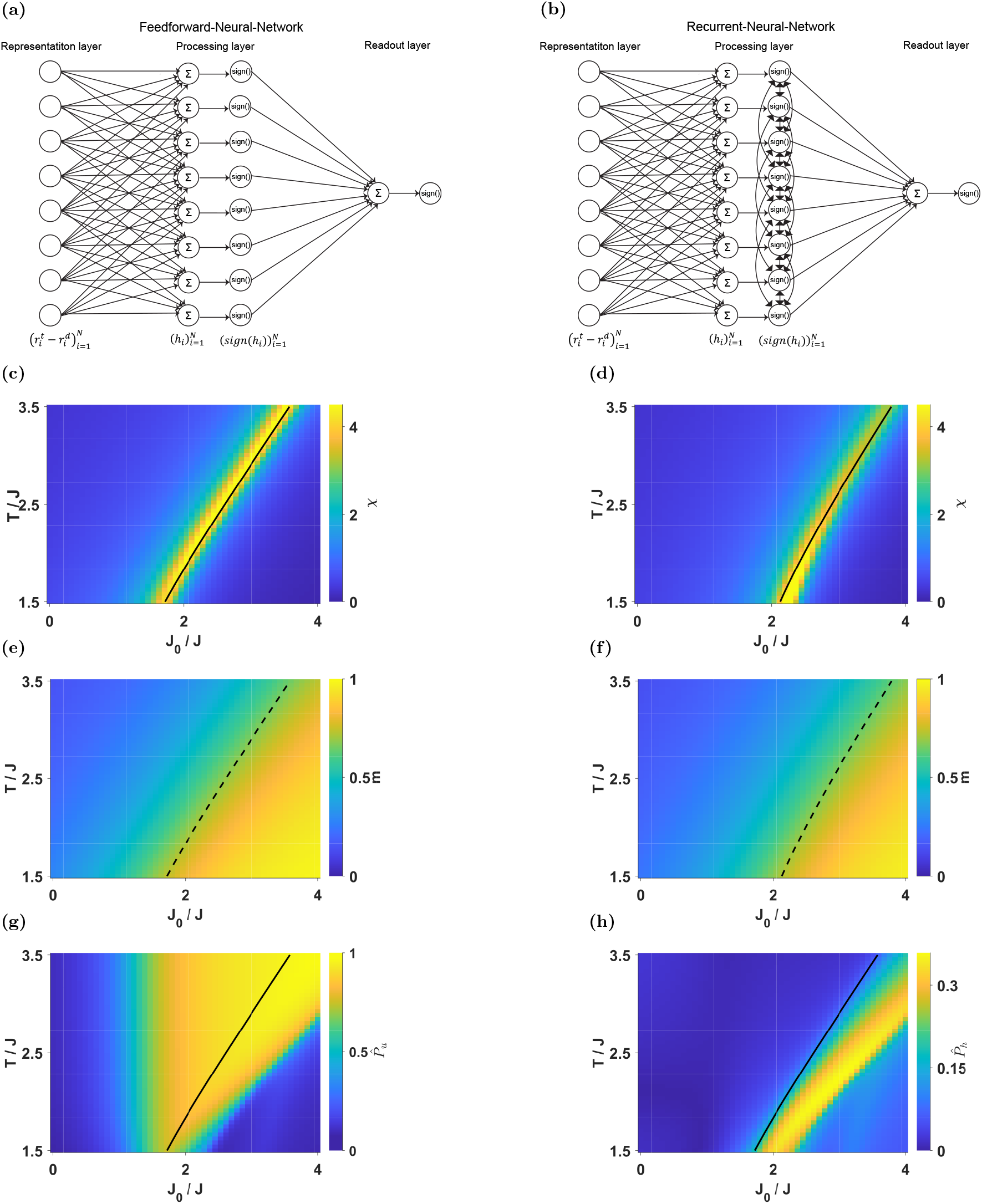
Coarse-tuning of the Feedforward Neural Network and the Recurrent Neural Network. **(a,b)** Schematic illustration of the (a) Feedforward Neural Network and (b) Recurrent Neural Network architectures. **(c**,**d)** Susceptibility in the *T/J* –*J*_0_*/J* parameter space for (c) Δ = 0.5 and (d) Δ = 1. The solid black line marks the phase transition. **(e**,**f)** Magnetization heat maps for (e) Δ = 0.5 and (f) Δ = 1. The dashed black line indicates the phase transition for the corresponding *h*_0_ = 0 case. **(g**,**h)** Projection of the signal onto (g) the uniform direction and (h) the random local field.

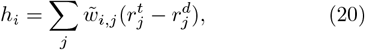

As above, we assume that the feedforward weight matrix, 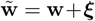, is a sum of two terms. The structured component, **w**, is identical for all neurons, 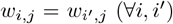. The second term, ***ξ***, is a random matrix representing the quenched noise of the synaptic weights. The elements {*ξ*_*i,j*_} are assumed to be i.i.d. Gaussian random variables with zero mean and 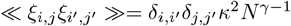.

Given the synaptic weights, **w**, the local fields, {*h*_*i*_},are random variables with a mean over trials 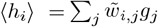. We denote the average taken over both trials and realizations by [·], yielding [*h*_*i*_] = *σ*_*g*_ ≡ *h*_0_. The covariance, 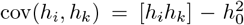, is given by (see Appendix E)

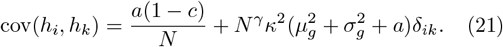

The first term on the right-hand side of Eq. 21 arises from the fact that all neurons in the processing layer receive identical structured inputs. This term represents the residual noise present in the optimal readout even under perfect fine-tuning of the weights.

In what follows, we neglect this contribution and approximate the distribution of the random fields, {*h*_*i*_}, by independent Gaussian variables with

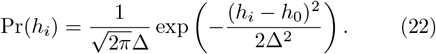

where,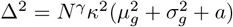.

Under the approximation of Eq. 22 the random fields, {*h*_*i*_}, are i.i.d. random variables with a variance scaling as *N*^*γ*^. This variability in the random fields induces heterogeneity in the responses of the neurons in the processing layer, {*s*_*i*_}. In the weak to moderate coarse-tuning regimes, −1 ≤*γ* < 0 this heterogeneity vanishes as the population size, *N* grows. We therefore focus on the strong coarse-tuning regime, *γ* = 0, where heterogeneity in both the random fields and the neuronal responses remains 𝒪(*N*^0^). Note that in the above approximation, the trial-to-trial fluctuations of responses of neurons in the processing layer, {*s*_*i*_}, are uncorrelated. Hence, we set *γ* = 0.

To estimate the stimulus from the neural responses, {*s*_*i*_}, we need to de fine a readout. A natural choice is a linear readout: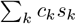, where *c*_*k*_ are the optimal {*s*_*i*_}, we need to de fine a readout. A natural choice weights for the processing layer. Consistency requires us to assume strong coarse-tuning in the readout weights from the processing layer as well. Consequently, adding feedforward layers cannot overcome the limiting effect of strong coarse-tuning. Below we consider the addition of recurrent connections to the processing layer.

### B. Recurrent Neural Network

Can recurrent connectivity overcome the quenched noise in the case of strong coarse-tuning? To address this question we incorporate recurrent interactions between neurons in the processing layer. Thus, we consider the network to comprise three layers: (i) a representation layer, 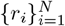, (ii) a single intermediate processing layer with neurons,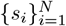, which are recurrently connected,and (iii) a decision unit denoted by 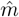, Fig.4b.

The representation layer 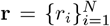 encodes the binary stimulus as previously, and has feedforward connections to the processing layer. Each processing neuron receives an i.i.d. quenched random local field {*h*_*i*_}, distributed according to Equation (22), which reflects the input from **r**. Crucially, unlike the purely feedforward architecture, the processing neurons could now interact via recurrent connections.

Each neuron in the processing layer is modeled as an Ising spin, *s*_*i*_ ∈ {±1} for *i* ∈ {1 … *N*s}. The network evolves according to Glauber dynamics at temperature 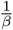, governed by the Hamiltonian:

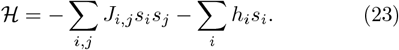

The recurrent connections {*J*_*i,j*_}, are taken to be i.i.d. quenched Gaussian random variables with:

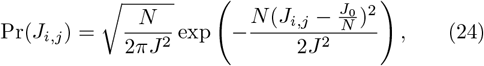

Where 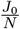 is the mean strength of the recurrent connec-tions and 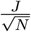is the standard deviation of the recurrent connections. The parameter *J* quenched disorder in the recurrent connections, similar to the strong coarse-tuning of the feedforward connectivity with *γ* = 0. Thus, the stochastic response of the Recurrent Neural Network neurons is naturally mapped to the Sherrington-Kirkpatrick (SK) model [42] with random fields [46, 47]. For simplicity, we assume that the trial-to-trial (thermal) fluctuations are more pronounced than the quenched noise, *T > J*. In this regime, one of the central order parameters characterizing the system represents strong is its magnetization (per spin),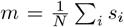. This system has been studied extensively, and in our regime of interest its phase diagram (for *h*_0_ = 0) is characterized by an ordered (ferromagnetic), *m >* 0, and unordered (paramagnetic), *m* = 0, phase, see [46–49] and appendix G.

When the recurrent connectivity exhibits a ferromagnetic bias, *J*_0_ *>* 0, the Recurrent Neural Network amplifies the signal in the structured component, *h*_0_, of the input fields. Figs. 4c & 4d illustrate this susceptibility, which quantifies the change in the Recurrent Neural Network response, *m*, to small changes in the input signal, 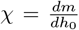. As shown, the susceptibility is maximal near the phase transition (marked by the black line). Comparing Figs. 4c & 4d, the main effect of the quenched disorder in the random fields, Δ, is that to obtain high signal amplification for larger values of Δ a stronger ferromagnetic bias, *J*_0_, is required. In the ordered phase, the structured component of the input fields acts as an ordering field that chooses the direction of the spontaneous magnetization, Figs. 4e & 4f. As in the case of susceptibility, larger values of Δ require a stronger ferromagnetic bias, *J*_0_, to obtain the same level of magnetization.

In the regions of the phase diagram in which the amplification is large, most of the signal resides in a two dimensional subspace spanned by the input fields, *h*, and the uniform direction, Figs. 4g & 4h. This takes us back to our initial problem: extracting the signal requires some degree of tuning of the readout weights.

## VIII. DISCUSSION

Here we investigated how coarse-tuning constrains the accuracy of population codes. We focused on the scaling of the signal-to-noise ratio (*SNR*) with population size for two readout algorithms: a naïve readout and an optimal readout. We found that the *SNR* of the naïve readout was largely insensitive to coarse-tuning, since its performance is already limited by noise correlations in the neural responses. In contrast, the *SNR* of the optimal readout exhibited distinct scaling regimes: it grew linearly with population size for weak coarse-tuning, crossed over to sublinear growth at intermediate coarse-tuning, and ultimately saturated for strong coarse-tuning.

Consequently, in the strong coarse-tuning regime, the *SNR* of both readout algorithms converged to finite limits, albeit for different underlying reasons, for large *N*. The asymptotic *SNR* of the optimal readout was governed by the magnitude of tuning fluctuations, quantified by *κ*, whereas the asymptotic performance of the naïve readout was dominated by noise correlations in the population activity. Finally, we showed that this fundamental limitation cannot be alleviated by adding further processing layers or by introducing recurrent connectivity.

How can our theory account for the high levels of accuracy observed in neural systems? One possibility is that the central nervous system operates in the weak to moderate coarse-tuning regimes, in which arbitrarily high accuracy can, in principle, be attained by an optimal readout given a sufficiently large population. However, we showed that in these regimes, the intrinsic heterogeneity of neuronal responses vanishes. This conflicts with extensive experimental evidence of persistent response heterogeneity [50, 51], which cast doubt on the hypothesis that neural circuits operate predominantly in the weak to moderate coarse-tuning regimes.

Alternatively, high decoding accuracy may be achieved in the strong coarse-tuning regime provided that the asymptotic *SNR* is sufficiently large. In this regime, the saturation of the *SNR* implies that pooling information beyond an effective population size, *N*_eff_, yields little additional benefits. Hence, the readout is robust to neuronal loss as long as the remaining population exceeds *N*_eff_ neurons.

Synaptic weights are inherently volatile, even in the absence of neuronal activity [26–28]. As a result, noise in synaptic strengths is expected to accumulate over time. A growing body of experimental evidence indicates that the response statistics of the representation layer can also evolve over time, a phenomenon commonly referred to as *representational drift* [52–56]. Together, these sources of variability are expected to degrade the performance of any fixed readout over time. Ongoing learning can partially counteract this degradation by continuously adjusting synaptic weights toward their target values. The interplay between stochastic synaptic fluctuations and corrective learning dynamics should thus produce a steadystate distribution of synaptic weights centered around their fine-tuned values. In the present work, we did not explicitly model the stochastic dynamics of synaptic plasticity; instead, quenched disorder was used to represent this steady-state distribution. A detailed investigation of the coupled dynamics of synaptic volatility and learning is beyond the scope of this study and is left for future work. Nevertheless, we believe that the present framework provides a foundational step in this direction.

With respect to learning, it is important to note that the naïve and optimal readouts make markedly different assumptions concerning the capabilities of the central nervous system. Learning the optimal readout generally requires a supervised learning mechanism, whereas the naïve readout can be acquired through comparatively simple forms of plasticity, such as homeostatic regulation. In particular, in our model, any homeostatic plasticity rule that induces independent fluctuations of synaptic weights around a common nonzero mean is sufficient to implement the naïve readout.

It is often assumed that the central nervous system operates near optimality [57]. However, because the performance of the naïve readout is comparable to that of the optimal readout and because the naïve readout can be implemented through substantially simpler learning mechanisms, the naïve readout remains a viable alternative. This reasoning is supported by two other potentially counterintuitive considerations.

The first is the invariant-manifold hypothesis which posits that the brain maintains functional stability despite ongoing synaptic volatility because the set of synaptic configurations that support a given computation forms a manifold (or low-dimensional subspace). Synaptic fluctuations are therefore substantial along this manifold, because they are strongly suppressed in directions orthogonal to it, thereby preserving function [33, 36, 53, 58–63]. In the context of our work, functionality was measured as the ability to extract the signal from the neuronal responses and estimate the stimulus. The signal was embedded by the response selectivity vector, **g** defined in Eq. 5. This vector can be decomposed into a fixed uniform component and zero-mean quenched fluctuations. The uniform direction, corresponding to the naïve readout, therefore spans the invariant manifold.

The second consideration arises from comparisons between theoretical readout performance and psychophysical measurements. Previous work has suggested that the naïve readout may be more consistent with empirical observations than the optimal readout [14]. Mendelson and Shamir [64] estimated the asymptotic accuracy of a naïve decoder (specifically, the population vector for angular estimation) and found performance levels comparable to those observed in psycho-physical experiments [65, 66]. Taken together, these observations raise the possibility that naïve readout mechanisms may play a role in neural decoding.

## ACKNOWLEDGMENTS

This work was supported by the Israel Science Foundation (ISF) under Grant No. 624/22 and Grant No. 824/21. The authors declare no competing interests.

## Appendix A

***SNR*^2^ of the fine-tuned naïve decoder**

Substituting the naïve choice of weights, 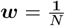, into Eq. 9, yields

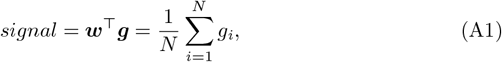

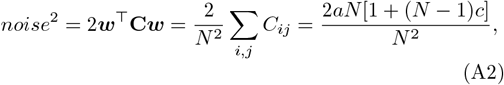

where we use in Eq. A2 the correlation matrix of Eq. 2. The *signal* is a random variable because it depends on the specific realization of the neuronal response selectivity, {*g*_*i*_}. Since the *g*_*i*’_s are i.i.d. random variables the *signal* is a self-averaging quantity with *signal* 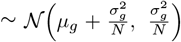 ; thus, in the limit of large *N* we can ignore the quenched variability of the *signal* and take it to be deterministic with *signal* = *µ*_*g*_. From Eq. 2, the *noise*^2^ has no quenched fluctuations and is 𝒪 (*N*^0^). Thus, the *SNR*^2^ of the naïve readout is

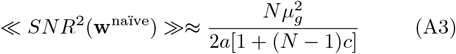

## Appendix B

***SNR*^2^ of fine-tuned Optimal decoder**

Substituting the optimal weights, ***w***^opt^ = **C**^−1^***g***, into Eq. 9, yields

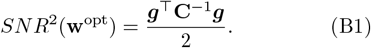

For our choice of correlation structure, Eq. A2, its inverse is given by

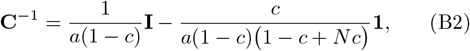

where **I** is the identity matrix and **1** is an all ones matrix. Thus, the *SNR*^2^ is a quenched random variable:

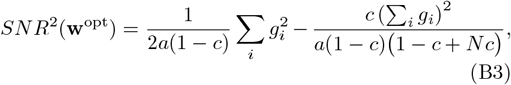

with mean

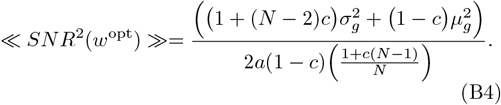

In the limit of large *N, SNR*^2^(*w*^*opt*^) ≈ *A*_1_ + *A*_2_, where 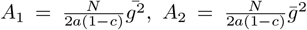and 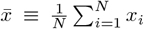 denotes spatial averaging of *x*. As both *A*_1_ and *A*_2_ are self averaging, so is *SNR*^2^(*w*^*opt*^).

## Appendix C

**The effect of coarse-tuning on the naïve decoder**

The *signal, s*, of the coarse-tuned naïve decoder, 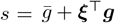, is a random variable with quenched mean of

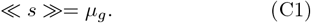

Denoting the quenched fluctuations of the *signal* by Δ*s* = *s*− ≪ *s* ≫, the quenched variance of the *signal*, ≪ (Δ*s*)^2^ ≫ is given by

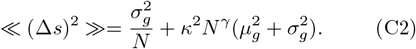

The squared *noise, n*^2^, of the naïve decoder is now a random variable

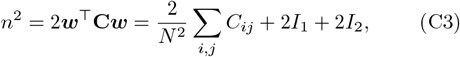

With

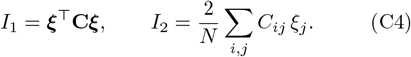

The quenched mean of the squared *noise* is given by

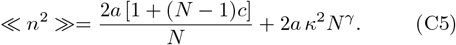

The quenched fluctuations of the squared *noise*, Δ*n*^2^ = *n*^2^ − ≪ *n*^2^ ≫, can be written as the sum of two terms: Δ*n*^2^ = Δ*I*_1_ + *I*_2_, where

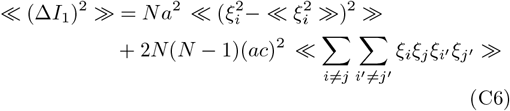

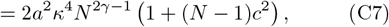

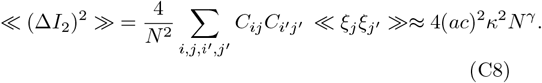

Thus, the quenched variance of *n*^2^ is

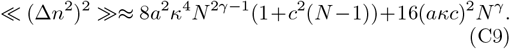

## Appendix D

**Optimal readout under coarse-tuning**

The quenched mean of the *signal* of the coarse-tuned optimal readout is

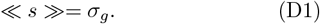

The quenched fluctuations of the *signal* are given by:

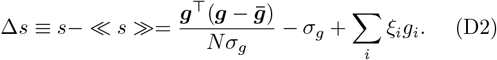

where, assuming large *N*, we approximated 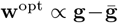 The quenched variance reads:

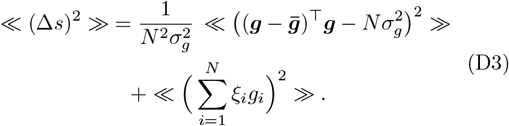

The contribution of the first term on the right hand side of Eq. D3 can be neglected for large *N*. Thus, the quenched variance of the *signal* is

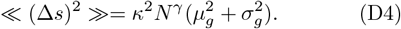

The squared *noise* is given by

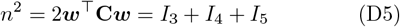

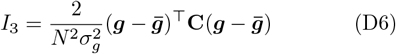

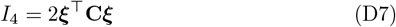

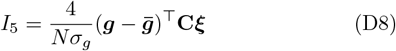

and its quenched mean is

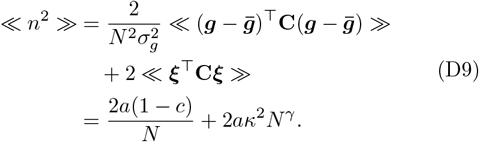

To compute the quenched variance of *n*^2^, note that *I*_3_ is the *noise* term of the fine-tuned optimal readout; hence, is self-averaging (see B). Now

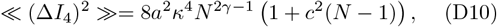

and

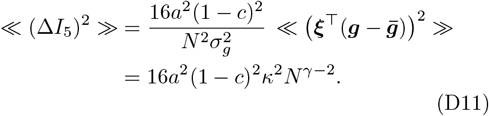

where we use in Eq. D11 the fact the 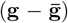 is an eigenvector of **C** with eigenvalue of *a*(1 − *c*). Thus, the quenched variance of *n*^2^ is given by

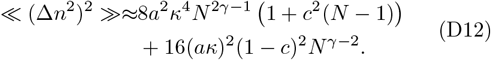

## Appendix E

**Feedforward neural network**

We consider a processing layer of *N* neurons with local fields

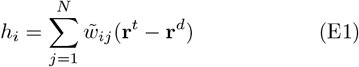

where,

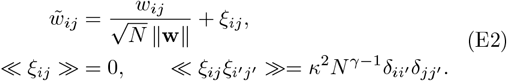

The mean of the local field is

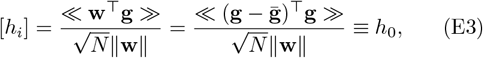

where [] denotes averaging with respect to both the trial-to-trial variability and the quenched disorder. The covariance of the local fields is given by

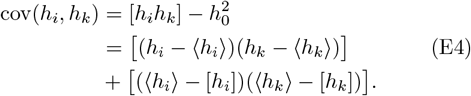

The first term on the right hand side of Eq. E4 yields

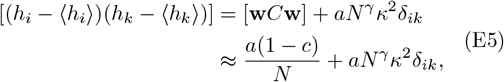

where for large *N* we approximated 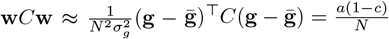. The second term yields

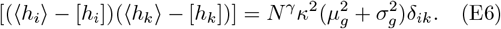

Thus, we obtain

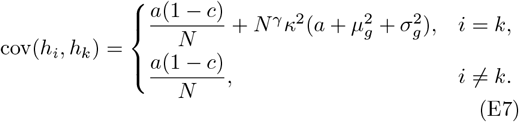

## Appendix F

**Sherrington-Kirkpatrick model**

The Recurrent Neural Network model of section VII B naturally maps to the Sherrington-Kirkpatrick model with random fields. This model has been extensively studied; see e.g., [42, 46, 47, 67]. For completeness we summarize the central results that are relevant to the current study.

We consider a system of *N* Ising spins, 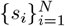, governed by the Hamiltonian in Eq. 23 at thermal equilibrium with inverse temperature *β*. The fields, {*h*_*i*_ }, are i.i.d. quenched Gaussian random variables, with the probability density

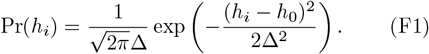

The recurrent connections, {*J*_*i,j*_ }, are i.i.d. quenched Gaussian random variables with

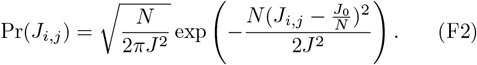

To appreciate the effect of recurrent connectivity it is insightful to first consider the case of *J* = 0. At thermal equilibrium the local magnetization, *m*_*i*_ = *s*_*i*_, is given by:

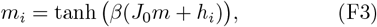

where *m* is the mean magnetization,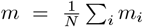which serves as an order parameter of the system. Thus, the self-consistent equation for *m* is

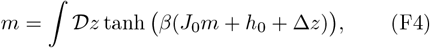

Where 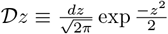 is the standard Gaussian measure. In the absence of external fields, *h*_0_ = 0 and Δ = 0, the paramagnetic state, *m* = 0, is always a solution of Eq. (F4). When the recurrent connectivity, *J*_0_, is stronger than some critical value, 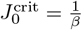the paramagnetic solution loses its stability and the system settles into one of the two solutions of the ferromagnetic phase. In this case, even a small ordering field, |*h*_0_| ≪ 1, will choose the sign of the ferromagnetic solution. Thus, in this case, the recurrent connectivity offers amplification of its input.

To understand the effect of random fields, Δ *>* 0, we first examine the case *J* = 0 and *h*_0_ = 0. In this case, due to the symmetry of Eq. (F4), a solution with *m* = 0 always exists. However, since Δ *>* 0 the local magnetization cannot be zero. Thus, the system is characterized by an additional order parameter:

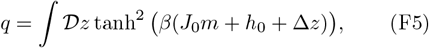

the Edward Anderson order parameter [67].

To find the the ferromagnetic transition (see [49]), one sets the mean of the external field to zero, *h*_0_ = 0, and, assuming a second order phase transition, expands Eq. F4 in small *m*. Using

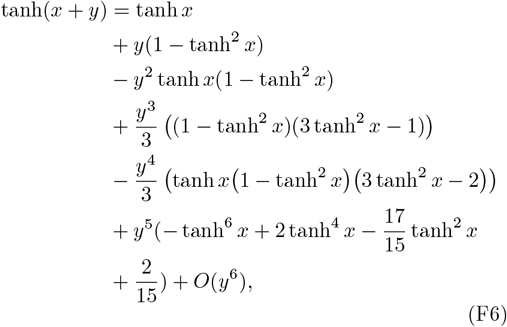

one obtains

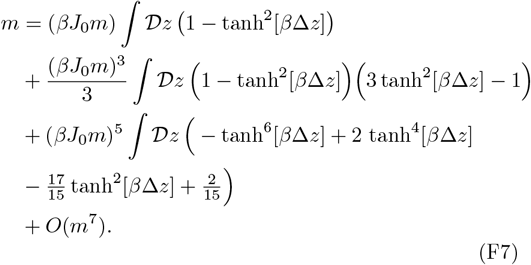

Equivalently, denoting *q*_2*k*_ ≡ ∫ 𝒟 *z* tanh^2*k*^(*β*Δ*z*),

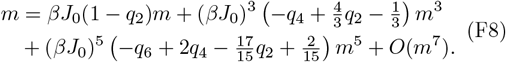

A second-order phase transition to the ferromagnetic phase occurs when the coefficient of the linear term in *m* in Eq. F8 changes sign

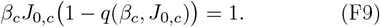

Note that under the assumption of Gaussian random fields, the coefficient of the cubic term in *m* in Eq. F8 is always positive; see [68]. Accordingly, a transition to the ferromagnetic phase can always be induced for sufficiently large *J*_0_. However, in the presence of random fields (Δ *>* 0), the transition is shifted to larger values of the recurrent coupling *J*_0_. Thus, fluctuations in the local fields (Δ *>* 0) attenuate the amplification of the ordering field *h*_0_ by the recurrent interactions.

In the general case of strong quenched disorder in the recurrent connections, *J >* 0, one can derive equations for the local magnetization that generalizes Eq. (F3) (see Appendix G):

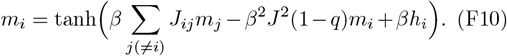

Within our chosen parameter regime, *βJ* < 1, the two order parameters *m* and *q* suffice to characterize the system [67]. Averaging Eq. (F10) over space and replacing the spatial average with averaging over the quenched disorder yields the self-consistent equations for the order parameters:

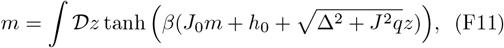

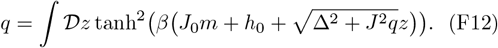

Similar to random fields (Δ *>* 0), the coupling of quenched disorder in the recurrent connections (*J >* 0) with nonzero *q*, suppresses amplification of the ordering field, *h*_0_. As a result, the transition to the ferromagnetic phase is shifted to larger values of the ferromagnetic bias *J*_0_. Nevertheless, for sufficiently large *J*_0_, the recurrent interactions still provide amplification of the signal, *h*_0_. We further map the Almeida-Thouless (AT) line [48], writing

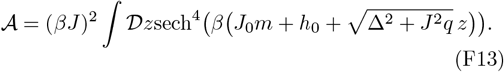

For 𝒜 < 1, the two order parameters in (F11) and (F12) fully characterize the system. The AT boundary is given by = 1; beyond it (𝒜*>* 1) the solution above is insufficient and additional techniques need to be utilized to characterize the system such as replica symmetry breaking [69–71].

## Appendix G

**TAP equation**

Here we derive TAP equations [72] for the Sherrington-Kirkpatrick model with local random fields using the cavity method [73]. The derivation presented below extends the method of Ref. [74] to include random local fields. The extension is straightforward and is presented here for completeness. The derivation proceeds in three steps.

### Step 1: adding a spin

We first extend the *N* -spin system by adding a single spin, *s*_0_, at site 0, which interacts with the existing spins via couplings 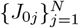drawn from the distribution in Eq. F2. The Hamiltonian of the extended (*N* + 1)-spin system is given by

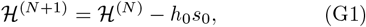

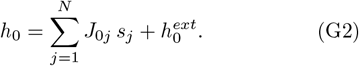

where *h*_0_ is the local field at site 0. The states of the extended system are distributed according to the Gibbs distribution with Hamiltonian ℋ ^(*N* +1)^:

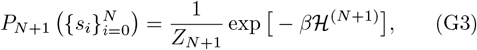

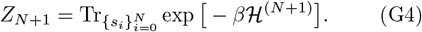

From Eq. G3 we obtain the marginal joint distribution of the cavity spin, *s*_0_, and the cavity field, *h*_0_,

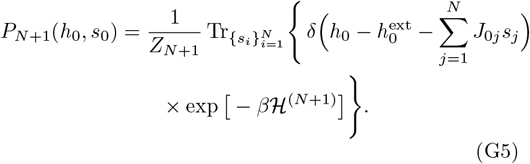

Using Eq. G1 we can write

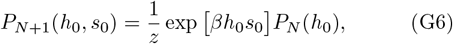

Where

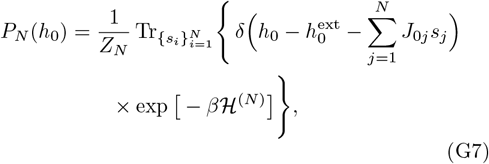

and the normalization, *z*, is given by

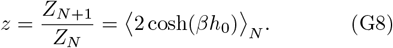

with⟨·⟩ _*N*_ denoting thermal averaging with respect to the *N* -spin system.

Averaging *s*_0_ and *h*_0_ with *P*_*N*+1_(*h*_0_, *s*_0_) gives the standard identities

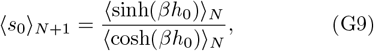

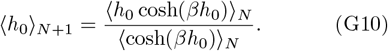

### Step 2: statistics of the cavity field *h*_0_

Next, we characterize the first two moments of the cavity field, *h*_0_, in the *N* -spin system:

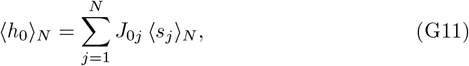

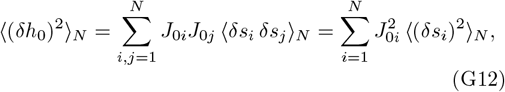

with *δX* = *X* −⟨*X*⟩_*N*_. In the last equality of Eq. G12 we used the fact that ⟨*δs*_*i*_*δs*_*j*_⟩ _*N*_ is of order 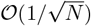and is independent of *J*_0*j*_, to approximate the sum over *i* and *j* only by the contribution of the *i* = *j* terms.

Defining 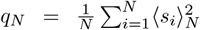 and using the selfaveraging of the right-hand side of Eq. G12 we obtain ⟨ (*δh*_0_)^2^⟩_*N*_ = *J* ^2^ (1−*q*_*N*_). Assuming the distribution of the cavity field, *h*_0_, in the *N* spin system is well-approximated by Gaussian distribution as the sum of *N* independent variables. Further assuming that *q*_*N*_ → *q* as *N* → ∞, the distribution of the cavity field is given by:

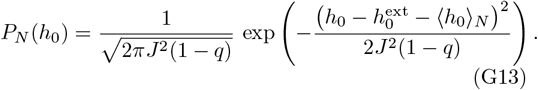

### Step 3: obtaining the TAP equation

Substituting (G13) into the Eqs. G9 & G10 yields

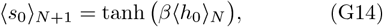

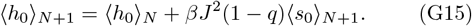

Substituting ⟨*h*_0_⟩_*N*_ from Eq. G15 into Eq. G14 and replacing site 0 with a generic site *i*, one obtains the TAP equations:

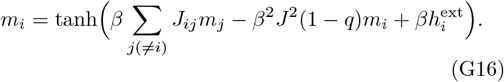

Note that, in this case 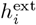 also exhibits quenched statis-tics.

#### The susceptibility matrix

The susceptibility matrix is defined by *χ*_*ik*_ ≡ *∂m*_*i*_*/∂h*_*k*_. The diagonal element of the susceptibility matrix can be obtained using the fluctuation-dissipation theorem, *χ* = *β***C**, where **C** is the correlation matrix *C*_*ij*_ = ⟨*δs*_*i*_*δs*_*j*_⟩. Since 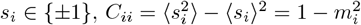. Thus,

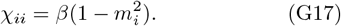

For the off-diagonal elements, we differentiate Eq. G16 with respect to the local field at site *k*, yielding

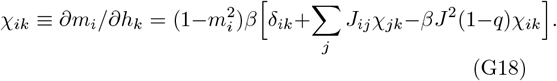

Defining the matrix **D** to be

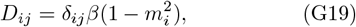

we obtain

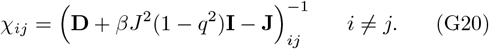

##### Figures 1-3: Neural population readout

Table II summarizes the parameter values used for the results in figures 1-3. Parameters marked with an asterisk (^∗^) vary across figure panels and are specified explicitly in the corresponding captions.

**TABLE 1:**
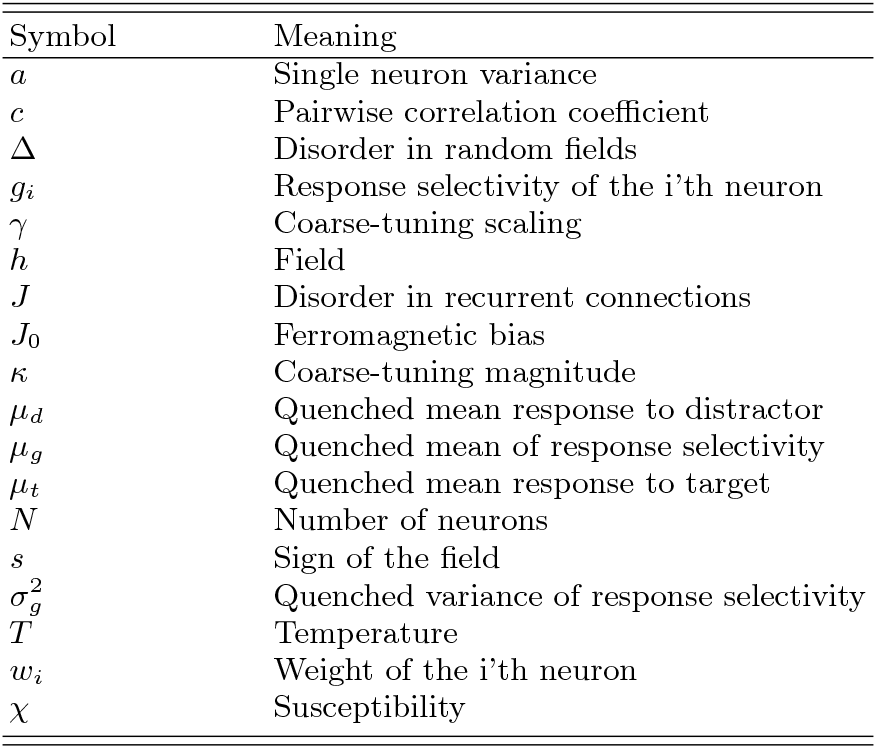
Alphabetical glossary of mathematical symbols used throughout the manuscript.

**TABLE 2:** Parameters for the neural population readout model (Figs. 1–3).

| Parameter | Description | Value |
| --- | --- | --- |
| $N$ | Number of Neurons | * |
| $a$ | Single-neuron variance | 12 |
| $c$ | Pairwise correlation coefficient | * |
| $\mu_t, \mu_d$ | Quenched mean response (t,d) | 12, 9 |
| $\mu_g$ | Quenched mean of response selectivity | 3 |
| $\sigma_g^2$ | Quenched variance of response selectivity | 24 |
| $\kappa$ | Coarse-tuning magnitude | * |
| $\gamma$ | Coarse-tuning scaling | * |

We performed 500 realizations of the quenched disorder. In each realization, we sampled 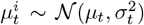 and 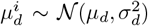 for each neuron *i*, yielding the se-lectivity 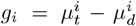 Coarse-tuning was drawn as *ξ*_*i*_∼ *𝒩*(0, *κ*^2^*N*^*γ*+1^). For each realization, we constructed the readout weights for the naïve and optimal decoders and computed the *SNR*. Finally we averaged the *SNR* over all realizations. The simulation results of the *SNR* (markers) were compared to the analytical expressions of the *SNR* (solid lines) derived in the main text.

##### Figure 4: Feedforward Neural Network and Recurrent Neural Network simulations

Table III summarizes the parameter values used for the Feedforward Neural Network and Recurrent Neural Network simulations. Parameters marked with an asterisk (^∗^) vary across figure panels and are specified explicitly in the corresponding captions. The coupling matrix and local fields were constructed according to Eqs. (22) and (24), respectively.

**TABLE 3:** Parameters for the Feedforward Neural Network and Recurrent Neural Network simulations (Fig. 4).

| Parameter | Description | Value |
| --- | --- | --- |
| $N$ | Number of neurons in processing layer | 200 |
| $J$ | Disorder in the recurrent connections | 1 |
| $h_0$ | Signal | * |
| $\Delta$ | Disorder in the random fields | * |
| $T$ | Temperature | [1.5, 3.5] |
| $J_0$ | Ferromagnetic bias | [0, 4] |

The TAP equations, Eq. (G16), were solved using damped fixed-point iteration with mixing parameter *α* = 0.35. Convergence was declared when 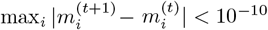, with a maximum of 4000 iterations.

The susceptibility matrix *χ*, Eq. (G18), was computed from the converged TAP solution. The dominant eigenvector of *χ*, **v**_max_ was obtained using MATLAB’s eigs function with tolerance 10^−10^; if this fails to converge, a full eigendecomposition was performed instead.

To analyze the structure of the dominant mode, Fig. 4g & 4h, we projected the unit-normalized eigenvector 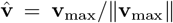 onto the subspace spanned by **û** = **1***/*∥ **1**∥ (uniform direction) and **ĥ**= **h***/*∥ **h** ∥ (normalized field). The projection coefficients (*c*_*u*_, *c*_*h*_) were obtained by solving

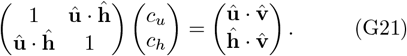

The residual 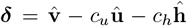 is orthogonal to both basis vectors.

The phase transition for *h*_0_ = 0, denoted in Fig. 4c-f by a black line, was computed as follows. For each temperature, we first solved Eq. (F12) for the Edwards-Anderson parameter *q*. The critical *J*_0_*/J* is then given by

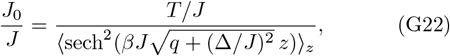

where *z ∼ 𝒩*(0, 1) and the expectation was evaluated using Gauss-Hermite quadrature with 80 nodes. All heat maps were computed on a 50 × 50 grid for a single disorder realization.

## Notes

### Competing Interest Statement

The authors have declared no competing interest.

### Summary of Updates

Figure 4 moved to result section.

